# Single nuclei multiomics reveals the drought-driven gene regulatory atlas in Arabidopsis

**DOI:** 10.1101/2024.01.11.575118

**Authors:** Jinbao Liu, Aqsa Majeed, Nilesh Kumar, Karolina M. Pajerowska-Mukhtar, M. Shahid Mukhtar

**Author notes:** These authors contributed equally to this work (Jinbao Liu, Aqsa Majeed).

## Abstract

The regulation of gene expression in plant responses to drought has been thoroughly investigated in previous studies. Despite this, a detailed understanding of the cell type-specific regulatory mechanisms, encompassing multi-layered biological processes, is lacking. In this study, we report the use of single-nucleus multiomic analysis in Arabidopsis seedlings in response to drought stress. Our single-nuclei RNA (snRNA) analysis delineated 14 distinct clusters representing major root and shoot cell types and discovered new cell type-specific drought markers. Integration of snRNA with single-nuclei ATAC (snATAC) data in leaf epidermis, root endodermis, and guard cells revealed accessible chromatin regions (ACRs)-linked genes predominantly enriched in pathways responsive to drought, heat, and light. Motif enrichment analysis and gene regulatory network (GRN) inference highlighted key transcription factors (TFs) and regulatory networks related to ethylene signaling pathways in endodermis as well as circadian rhythms in both endodermis and guard cells. Pseudotime analysis identified critical transcriptomic progression from metabolic process to stress response within three cell types. Overall, this study elucidates drought-related regulatory mechanisms in Arabidopsis at single-cell resolution, providing valuable insights into the fundamental regulatory events involved in stress responses. It also serves as a reference for future single-cell multiomic investigations in crop plants.

**One Sentence Summary:** Single cell multiomic analysis under drought stress

## Introduction

Drought, a recurring climatic phenomenon marked by prolonged periods of water deficit, poses significant challenges to ecosystems, economies, and public health (*1, 2*). For plants, the detrimental impacts of drought include impaired growth, delayed development, reduced reproduction, and heightened susceptibility to diseases, leading to lower yields and heightened mortality (*3-5*). Previous studies on elucidating plant-drought interactions have revealed multi-layered responses, encompassing genomics (*6-9*), epigenomics (*10-12*), transcriptomic (*10, 13, 14*), and proteomics (*15-17*). Omics-wide profiling has advanced our understanding of the complex and integrative regulation involving various biological molecules in plant responses to drought (*6-17*). Mounting evidence highlights the varied drought response exhibited by distinct plant cell types. For instance, in both rice and Arabidopsis, root endodermis has been shown to accumulate suberin in its cell walls as drought progresses, aiming to minimize water loss in roots (*18, 19*). In addition, abscisic acid, a major phytohormone in drought stress signaling, is observed to regulate distinct transcriptomic signatures in leaf epidermis and guard cells (*20*). However, these past studies on drought have primarily been conducted using bulk tissue or organs, often overlooking the specific contributions of individual cell types within tissues. As such, the mechanism by which distinct cell types coordinate to establish a unified and organized response to drought is yet to be fully elucidated.

The recent flurry of single cell studies has significantly advanced our understanding of cell-specific profiles in plant species (*10, 21-33*). The use of single-cell RNA sequencing (scRNA-seq) or snRNA-seq has shed light on the transcriptomic heterogeneity within plants, unveiling shared and distinctive signatures that define cell identities and functions (*21-29*). snATAC-seq has been instrumental in mapping the open chromatin landscapes of plants, aiding in the identification of the key regulatory factors involved in various cell types (*10, 30-34*). However, single-layered profiling often fails to provide the complete landscape of regulatory interactions. To address this knowledge gap, integrative analysis combing scRNA/snRNA-seq data with snATAC-seq has been employed across species (*35*) to unveil the cellular regulatory mechanisms (*10, 30, 33*). Concurrently, a particularly notable technical advancement are single-nucleus multiome (snMultiome) techniques; this innovation allows for the simultaneous profiling of snRNA and snATAC landscapes within the same cells. It has been shown promising in plant species such as Arabidopsis and cotton (*35, 36*). The robust potential of the multiome now opens new avenues for an in-depth understanding of the intricacies of cellular regulatory events in plant response to drought.

One key regulatory process in plant cells includes the modulation of target genes (TG) expression by TFs through interactions with regulatory elements, such as promoters and enhancers (*37-40*). Determination of TF binding site (TFBS) enrichment and inference of GRN are essential to decipher the intricate network exhibited by TFs and TGs (*38, 39, 41*). In a single-cell context, a suite of GRNs tools have been developed to harness scRNA/snRNA or snATAC data to delineate the core regulatory networks within specific cell types (*38, 39, 41-44*). In plants, two representative tools are MINI-EX (Motif-Informed Network Inference method based on single-cell EXpression data) and MINI-AC (Motif-Informed Network Inference Based on Accessible Chromatin) (*38, 39*). Both tools are compatible with different types of single cell dataset; MINI-EX infer networks based on scRNA or snRNA data whereas MINI-AC mainly focus on snATAC data (*38, 39*). These tools have been effectively applied to single cell data from plants including Arabidopsis and maize. Remarkably, MINI-AC has the capacity to integrate both scRNA/snRNA and snATAC data into a comprehensive analysis. This approach offers insights into cell-type-specific regulatory networks and facilitates the identification of differentially expressed TFs (*39*). These features highlights MINI-AC significance in probing more complex and integrative single cell datasets such as snMultiome (*39*).

To decipher the intricate drought-driven TF-TG regulatory network in a cell type-specific manner, we employed snMultiome analysis into 7-day-old Arabidopsis seedlings that were subjected to drought stress. Profiling entire seedlings enabled the capture of cellular similarities and differences arising from shoots and roots. snRNA analysis revealed significant transcriptomic changes in distinct cell types, along with identifying new cell type-specific markers under drought. Furthermore, by integrating snRNA-seq and snATAC-seq, we identified important TFs and TGs as well as mapped core regulatory networks in major cell types. Finally, trajectory analysis unveiled a sequential gene expression pattern of gene expression correlating with drought progression, particularly in the endodermis, epidermis, and guard cells. This research sheds light on the complex cell type-specific gene regulatory atlas involved in drought, emphasizing the significant roles of various cell types in adapting to drought conditions.

## Result

### Single-Nucleus Multiomic Atlas of 7-Day-Old Seedlings under Drought Stress

To evaluate the effect of drought on whole plants, we grew Columbia-0 (Col-0) seeds in half Murashige and Skoog (MS) media for 6 days (Figure 1A). Subsequently, half of the seedlings were subjected to a chemically induced drought stress by transferring them to polyethylene glycol (PEG)-infused ½ MS media for 16h drought stress (*45, 46*). The remaining seedlings served as controls (Figure 1A). To validate the efficacy of drought stress, we quantified the expression of four well-established drought-responsive genes (RD29A, RD29B, ERD1, and ABF3) using quantitative reverse transcription polymerase chain reaction (qRT-PCR) (Figure S1A). The significant upregulation of these marker genes under drought stress verified the activation of both abscisic acid (ABA)-dependent and independent pathways, two main drought stress signaling cascades (*3*). Following the treatment, samples were collected under both conditions, and we proceeded to isolate nuclei from the 7-day-old Arabidopsis seedlings using a gradient centrifugation approach (See Method). After nuclei staining and Fluorescence-Activated Nuclei Sorting (FANS), 16,000 nuclei per sample were loaded onto 10x Chromium Multiome platform (Figure 1A), with three replicates to each treatment. After sequencing, the snRNA-seq and snATAC-seq were preprocessed via CellRanger-ARC (v 2.0.2) and aligned to the Arabidopsis transcriptome and genome, respectively. A total of 15,376 transcriptomes from all samples met the stringent filtering criteria, which led to the identification of 24,504 genes covering approximately 90% of the total protein-coding genes in Arabidopsis. Owing to the incompatibility of the 10x CellRanger-ARC peak calling algorithm with Arabidopsis, we applied a customized approach that utilized nuclei filtered with RNA strandards to perform MACS2 peak calling to ATAC reads.

**Fig. 1:**
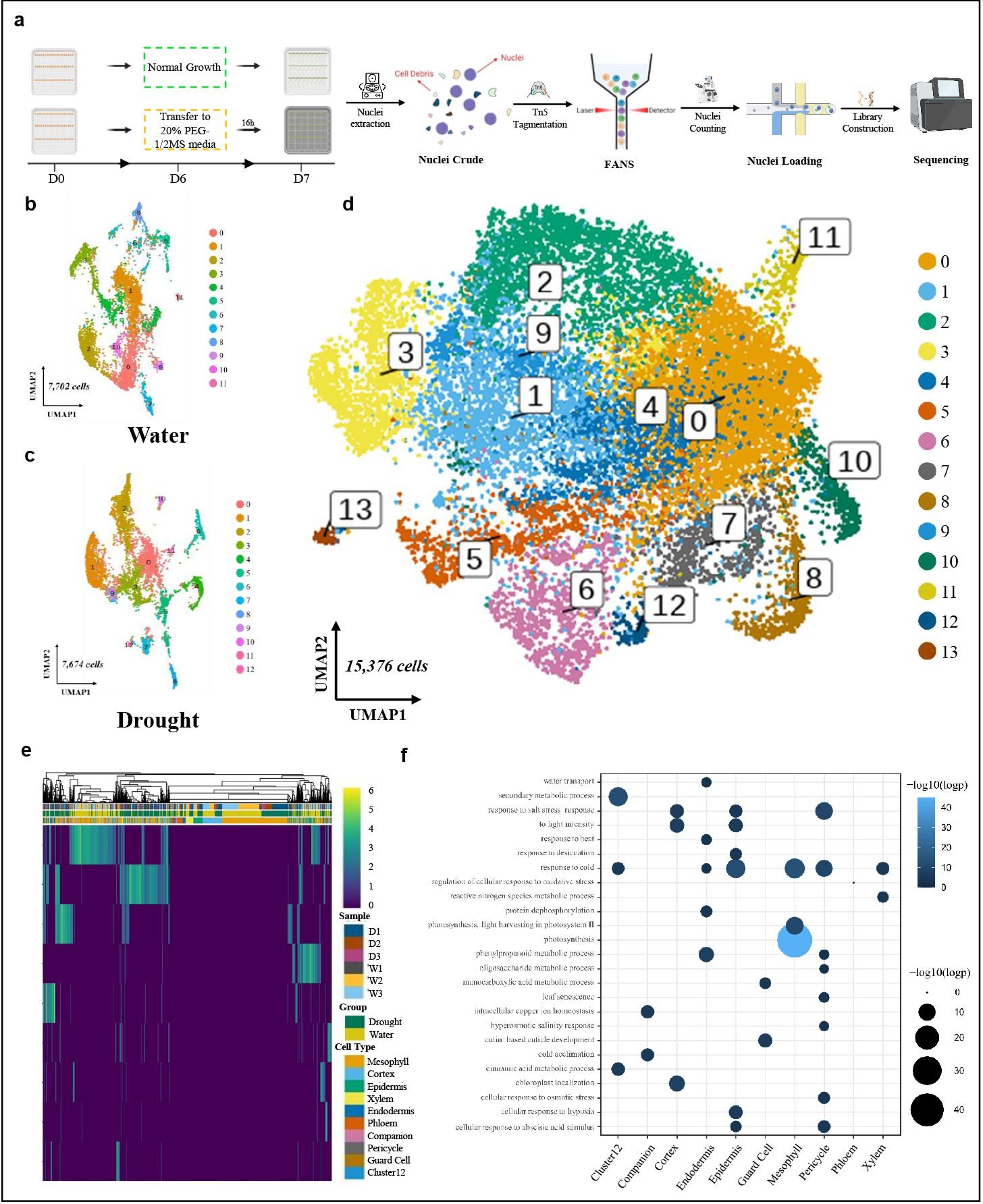
Single nuclei RNA analysis reveals shared and unique cell type-specific transcriptomic signatures under drought. (A) Schematic of drought simulation and single nuclei multiomics (B) Uniform Manifold Approximation and Projection (UMAP) plots of snRNA clusters under water conditions. Nuclei are colored by Leiden clusters. (C) UMAP plots of snRNA clusters under drought conditions. Nuclei are colored by Leiden clusters. (D) UMAP plots of snRNA clusters after embedding drought and water groups. Nuclei are colored by Leiden clusters. (E) Heatmap showing differential expressed genes (DEGs) exhibited by cell types. The sidebar shows sample groups, treatment group and cell types. (F) Gene set enrichment analysis of marker genes in each cell type.

To discern distinct cell types, we initially analyzed the snRNA-seq data using dimensionality reduction and clustering for both groups, namely drought and no treatment. The analysis unveiled similar numbers of distinct clusters, 12 and 13, respectively (Figure 1B and 1C). Through the integration of data from both groups using Seurat, we identified a total of 14 clusters. The number of nuclei in each cluster ranged from the highest in cluster 0 to the lowest in cluster 13 (Figure 1D and S1B). We assigned the major cell types to 13 out of 14 clusters by comparing cluster markers with a set of well-characterized cell type markers, with cluster 12 being unidentified (Figure 1D and S1C). Among identified clusters, mesophyll in clusters 0, 1, and 4 constituted the majority of the nuclei, followed by cortex, epidermis, xylem, endodermis, phloem, companion, pericycle, and guard cells (Figure S2A). The consistent cell type ratios across different replicates and treatments underscored the reproducibility of our nuclei isolation and analysis methods (Figure S2B). Notably, some cell markers exhibited condition-biased expression (Figure S2C). For example, epidermis markers AT2G15050 (LTP) and AT2G48130 (LTPG15) show water- and drought-related expression patterns, respectively (Figure S1C). It suggests that the expression variability of cell-type markers might influence the reliability of previously established markers in different conditions. In conclusion, the identified cell-type markers successfully delineated the major cell types present in Arabidopsis roots and shoots under drought conditions.

### Transcriptomic Analysis Revealed the Shared and Unique Drought-driven Cellular Response among Cell Types

To elucidate the cell type-specific transcriptomes under drought, we compared their transcriptome profiles between drought-stressed and control conditions in each cell identity. Our analysis identified a total of 11,019 differentially expressed genes (DEGs) that were either upregulated or downregulated in specific cell types (Figure 1E and S2C). Pseudobulk analysis yielded 12,215 DEGs, slightly higher than the total number of DEGs summed up from individual cell types. Specifically, we discovered 3,304 DEGs unique to single-cell analysis and 4,500 DEGs unique to pseudobulk analysis. This suggests that both analyses could reveal distinct transcriptomic aspects. Further comparison across cell types revealed that 3,983 genes were uniquely expressed in only one cell type, while 6,964 genes were shared by two or more cell types. Identifying these drought stress-related DEGs aids in discovering new, cell type-specific markers induced by drought. For instance, AT1G11720 (SS3), a starch synthase-encoding gene crucial for controlling starch biosynthesis (*47*), was uniquely upregulated in guard cells under drought. This upregulation implies a pivotal role for SS3 in drought-induced guard cell activities, given that starch accumulation in guard cells has been shown to increase osmotic stress (*48, 49*). Another gene, AT2G17290 (CPK6), encodes a calcium-dependent protein kinase 6, and was found to be specifically regulated in the pericycle under drought conditions. Given its role in phosphorylating ABA-responsive element binding factors and its cell-to-cell mobility (*50, 51*), the upregulation of CPK6 in pericycle might indicate a root-shoot interplay during drought. Overall, the analysis of DEGs has provided a comprehensive view of the unique and shared upregulated or downregulated gene signatures involved in different cell types during drought stress.

Furthermore, to investigate the potential function as a function of the cell type-specific gene expression, we initially performed a gene set enrichment analysis on cell type-specific marker genes (Figure 1F and S3 This revealed shared pathways including “response to cold,” “response to salt stress,” and “response to intensity” across multiple cell types, such as epidermis, guard cells, root endodermis. The unique pathways exhibited by cell types are consistent with their primary functions in tissues. For example, root endodermis cells demonstrated pathways enrichment in “water transport,” “response to heat,” “protein dephosphorylation,” and “phenylpropanoid metabolic process”; leaf epidermis cells exhibited pathways like “response to light intensity” and “cellular response to abscisic acid stimulus.” Remarkably, mesophyll cells showed enriched pathways related to photosynthesis, underscoring their critical role in sustaining photosynthesis metabolism under drought. Taken together, the pathway analysis highlights the complexity of plant cellular function in stress response.

### Linkage between snRNA and snATAC Identified Drought-Induced Gene Expression Regulated via Accessible Chromatin Regions (ACRs)

Our analysis using snATAC did not yield distinct separations between clusters, indicating potentially less inconsistency in chromatin accessibility between cells (Figure S4A). To decipher the cell type-specific ACRs with more accuracy, we integrated snRNA and snATAC data from both groups into a unified pattern, which shows a better resolution compared to snATAC clustering alone (Figure 2A, S4A and 4B). This integration by Seurat delineated consistent cell types compared to snRNA clustering (Figure 2A, S4A-C) and established overall snATAC-snRNA linkages across cell types. Linkage distribution along transcription start sites (TSSs) shows it most frequently happens within 250bp upstream and downstream of TSSs in both groups (Figure 2B, 2C, S4C-E). Notably, in-depth linkage analysis into overall cell types displayed the cell type-specific or condition-specific expression patterns of linked genes. For example, AT1G04680 is present in the drought-stressed epidermis dataset but is not linked to other cell types or control conditions (Figure S4F-H). Most cell type markers appear to be associated with specific cell types regardless of conditions. Taken together, this variation among linked genes suggests the ACR-based gene regulation generated by cell types or conditions.

**Fig. 2:**
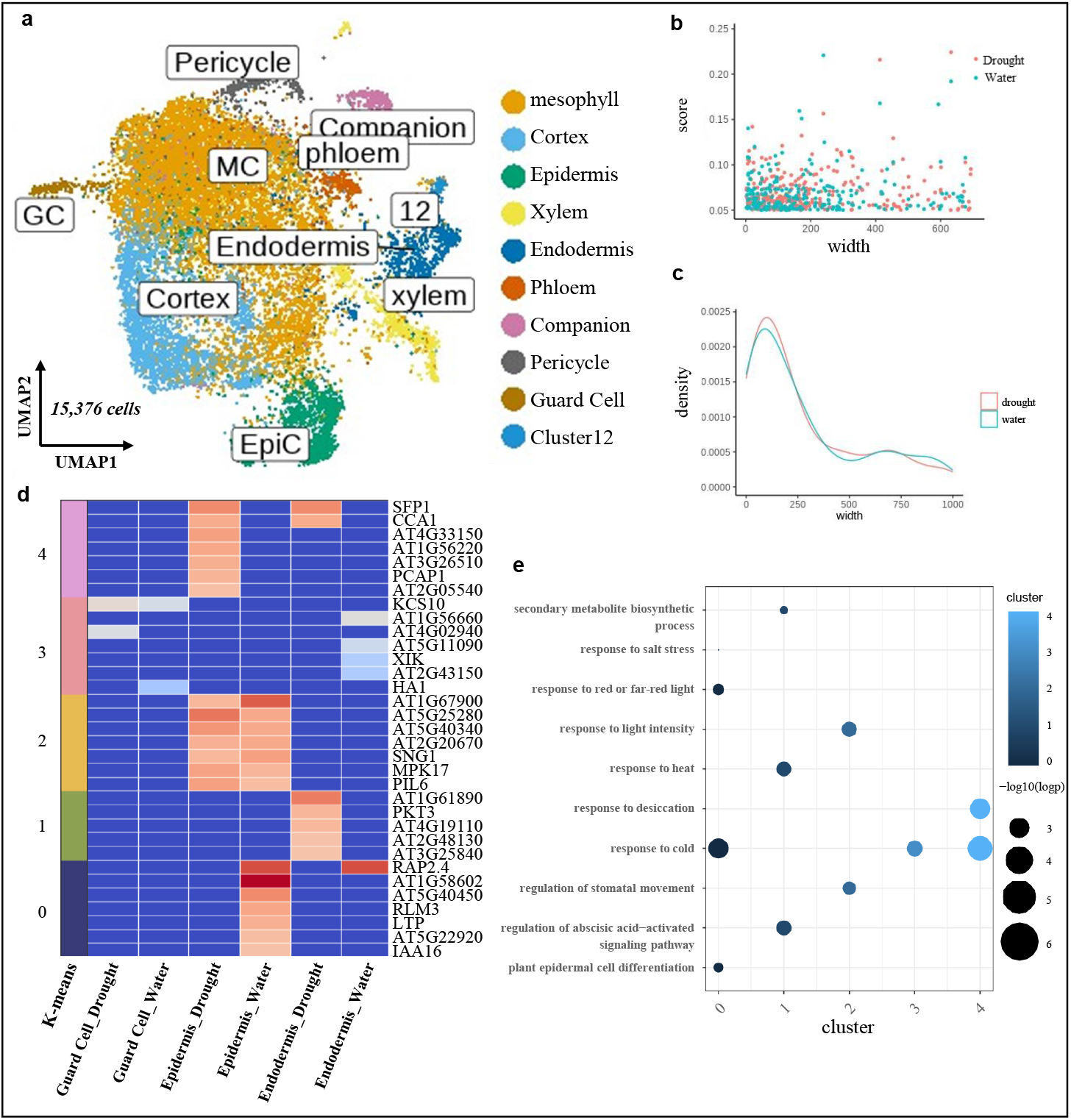
Linking transcriptomic profiles and chromatin accessibility identified cell type-specific and condition-specific regulatory events. (A) Uniform Manifold Approximation and Projection (UMAP) plots of embedding chromatin accessibility and gene expression landscapes from drought and water groups. Nuclei are colored by Leiden clusters. (B) The frequency of linkage between gene expression and chromatin accessibility at the width 5000bp of upstream and downstream of transcription start sites (TSS) for drought and water samples. (C) The frequency of linkage between gene expression and chromatin accessibility at the width 1000bp of upstream and downstream of transcription start sites for drought and water samples. (D) K-mean clusters heatmaps of linked genes among main cell types (root endodermis, leaf epidermis and guard cell) and conditions. The sidebar indicates score of k-mean clusters. (E) Gene Set Enrichment Analysis of linked genes among main cell types (root endodermis, leaf epidermis and guard cell)

To further explore the detailed function of linked genes, we selected endodermis, epidermis, and guard cells — all of which have been widely studied in previous drought studies (*18-20*) — for expression pattern visualization and functional analysis. *K*-means clustering identified sets of genes unique to conditions or cell types (Figure 2C). Compared to endodermis and epidermis, guard cells exhibited a less variable linked gene expression, likely due to the lower ATAC and RNA reads confined by limited nuclei. The functional analysis of linked genes highlighted involvement in multiple drought-responsive pathways, such as “response to desiccation” and “regulation of abscisic acid-activated signaling pathway”(Figure 2D). Interestingly, light signaling and heat-responsive pathways including “response to red or far-red light”, “response to light intensity”, and “response to heat”, are shown to be clustered among linked genes, which indicates plants could regulate light- or heat-related genes expression by interacting with open chromatin regions under drought stress (Figure 2D). Taken together, the linkage analysis indicates drought could regulate the gene expression via interacting with ACRs in a cell type-specific manner.

### Motif Enrichment Analysis and MINI-AC Infer Core TFs and TF-TG Regulatory Networks under Drought

To explore the crucial TFs involved in regulating drought response, we conducted motif enrichment analysis by using snATAC data after integration. This analysis identified a total of 447 transcription factors (TFs) across various cell types. Notably, a substantial number of TFs involved in ethylene signaling pathways are enriched in root endodermis, suggesting a critical role of ethylene signaling in endodermis response to drought. In addition, 195 out of 447 TFs were uniquely associated with a single cell type, while the remaining 252 were shared across two or more cell types. For example, guard cells uniquely expressed AT5G48560 (bHLH78), a TF known for its role in regulating flowering time, which was shown to be notably upregulated in shoots and roots during the later stages of drought stress (*52*). Additionally, endodermal cells distinctively expressed AT4G38620 (MYB4), a member of the R2R3-subfamily of TFs involved in cadmium stress tolerance and flavonoid biosynthesis (*53, 54*). Epidermal cells were characterized by AT1G49720 (ABF1), involved in the regulation of various stress-responsive genes (*55*), although its cell type-specific roles in drought response remain unclear. Moreover, out of the 252 shared TFs, 187 were exclusive to two cell types. Examples include AT3G23690 (bHLH77), AT5G05790, and AT3G52440, all predominantly expressed in guard cells and epidermal cells. This differential motif enrichment was instrumental in uncovering novel TF deployments in specific cell types which might be covered by bulk analysis (Figure 3A and S5A). For instance, ABF2, another key TF responsive to ABA signaling in addition to ABF1, exhibited pronounced binding site enrichment in bulk analysis but was less prominent in individual cell types.

**Fig. 3:**
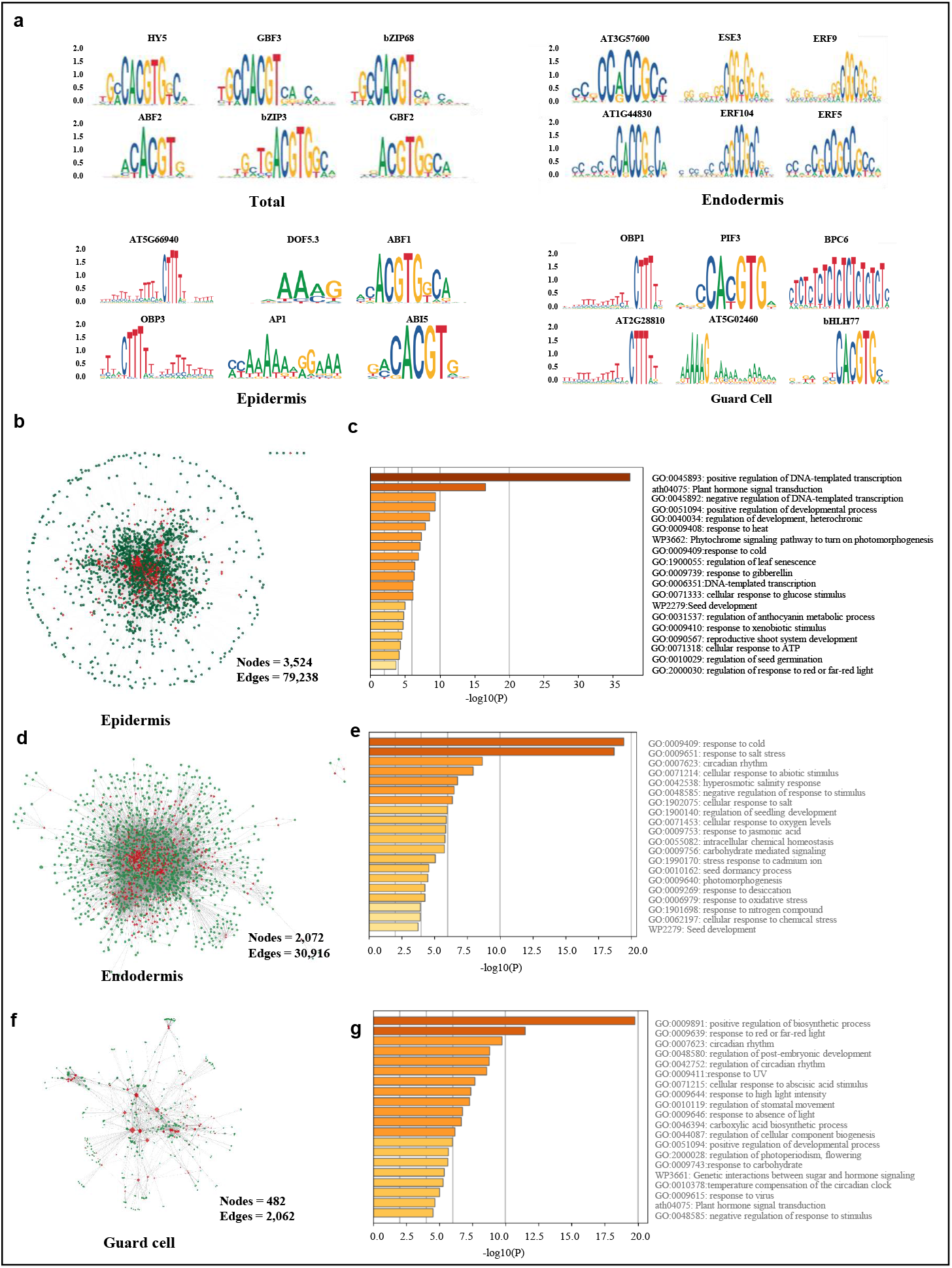
Network biology by integrating single nuclei RNA and ATAC data leads to find new, single cell-based core transcriptional factors and regulatory networks under drought. (A) Top motif enriched in whole seedling level and main cell types including epidermis, endodermis and guard cells. (B) Gene regulatory network inferred by MINI-AC in epidermis. (C) Pathway analysis of inferred gene regulatory network in epidermis. (D) Gene regulatory network inferred by MINI-AC in endodermis. (E) Pathway analysis of inferred gene regulatory network in endodermis. (F) Gene regulatory network inferred by MINI-AC in guard cell. (G) Pathway analysis of inferred gene regulatory network in guard cell.

Next, to further harness both snRNA and snATAC data to form drought-driven regulatory networks within cell types, we employed MINI-AC. It was previously demonstrated that MINI-AC is effective with snATAC data obtained from different leaf Arabidopsis and maize leaf tissues. Additionally, it allows for the combined analysis with DEGs derived from snRNA analysis (*39*). We partitioned our dataset by creating subsets for each cell type that included both ACRs and DEGs. With a small subset, we can infer the GRN and core codes presenting in each cell type. For instance, the inference reveals 3,524 nodes and 79,238 edges in the epidermis, 2,072 nodes and 30,916 edges in the endodermis and 482 nodes and 2,062 edges in guard cells (Figure 3B). Notably, the GRN in the endodermis, epidermis, and guard cells comprised 155, 263, and 35 hub-nodes, respectively. GRN pathway analysis indicated a predominance of stress-related transcription factors (TFs) in the epidermis and endodermis, implicated in pathways such as “response to cold” and “response to salt stress”. Conversely, TFs in guard cells were enriched for functions in “positive regulation of biosynthetic process”, “response to red or far-red light”, and “circadian rhythm”, suggesting a distinct regulatory network (Figure 3C, E and G).

Additionally, focusing on core TFs, we were able to further construct detailed TF-specific regulatory networks within cell types (Figure S5B, C, and D). For example, two significant epidermal markers, AT1G07640 (OBP2) and AT2G28510 (DOF2.1), formed networks with 852 nodes and 824 edges, and 822 nodes and 821 edges, respectively. Interestingly, these TFs did not exhibit significant up- or down-regulation in snRNA data, yet their prominence in the combined analysis underscores their vital roles in the epidermis. This implies that integrating both snATAC and snRNA datasets in GRN inference can uncover key regulators that might remain concealed by snRNA analysis alone.

### Pseudotime Analysis Reveals Gradual Gene Expression Changes in Main Cell Types

Utilizing snRNA data, we conducted an extensive pseudotime analysis across entire all cell types to unravel the developmental progression of various cell types (Figure 4A and S6). Specifically, beginning with the root endodermis, this trajectory revealed a gradient expression transition that progressed to the leaf epidermis and vice versa (Figure S6, S7A and B). The opposite pattern highlights a dynamic interplay between both cell types. Given the critical role of root endodermis, leaf epidermis, and guard cells discussed above, we concentrated our pseudotime analysis by subsetting these cells out of whole clusters (Figure 4B). The inferred trajectories, originating from the root endodermis, demonstrated a gradual shift in transcriptomic profiles, ultimately converging in guard cells (Figure 4B). Throughout this trajectory, we highlighted a multitude of genes uniquely expressed.

**Fig. 4:**
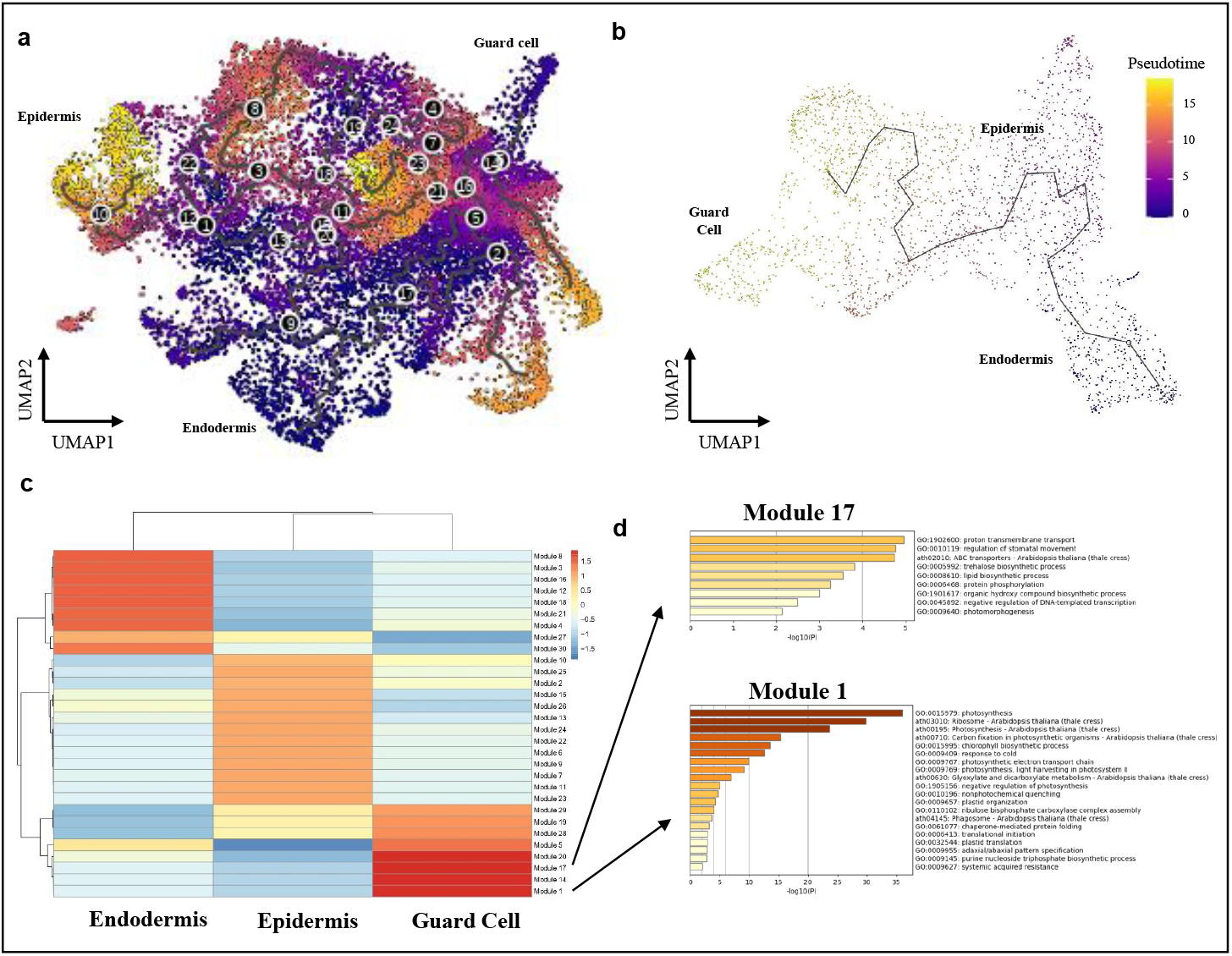
Pesudotime analysis of main cell type reveals a gradual transcriptomic transition along endodermis, epidermis and guard cell. (A) Inferred pseudotime trajectory by monocle3 in all cell types, with endodermis as start point. (B) Inferred pseudotime trajectory by monocle3 in subset of whole clusters, including endodermis, epidermis and guard cell. (C) Heatmap showing the grouped modules by variable gene expression along with pseudotime trajectories of endodermis, epidermis and guard cells. (D) Pathway analysis of modules inferred by monocle3, exemplified by module 17 and module 1.

Next, employing Monocle3 (v1.3.4) for exploring co-regulated modules, we systematically categorized variable genes into 30 distinct modules. Notably, while some were shared by cell types, most modules displayed a high degree of cell-type specificity (Figure 4C). For example, modules 1, 14, 17, and 20 showed significant enrichment in guard cells. Gene Ontology analysis of these modules indicated that modules 1 and 20 were related to photosynthesis pathways, including “photosynthesis”, “adaxial/abaxial pattern specification”, and “plastid organization”. On the other hand, modules 14 and 17 aligned more closely with stress-related pathways, such as “stomatal complex development”, “wax biosynthesis process”, and “response to UV”. Further analysis of these cell-type-specific modules unveiled a sequential function exertion across cell types (Figure S7C). Specifically, the gene expression within endodermis initiates metabolic process-related pathways and gradually shifts to drought-responsive pathways. In the epidermis, early stages are marked by enrichment in cell growth and hormone signaling pathways, while later stages predominantly exhibit stress-related modules. Intriguingly, in the initial stages of guard cell gene expression, we identified a cluster of “response to salicylic acid” genes, underscoring the potential significance of salicylic acid signaling pathways in drought stress response [66]. Taken together, this analysis reveals a gradual transcriptomic progress, which exerts distinct functions along with pseudotime trajectory of three cell types.

## Conclusion

In conclusion, our study, through the construction of a single cell multiomic atlas of Arabidopsis seedlings, provides novel insights into the plant’s cellular response under drought stress. The distinct transcriptomic profiles and accessible chromatin patterns identified across various cell types have unveiled the asymmetric nature of gene regulation shaped by regulatory elements and expressed genes. Importantly, the integration of snRNA and snATAC data to infer GRN has laid the foundation for understanding the intricate network within main cell types such as root endodermis, leaf epidermis, and guard cells. Furthermore, the trajectory analysis of drought progression in three cell types has not only confirmed the gene expression dynamics over time course but also highlighted the involvement of distinct pathways, such as salicylic acid. This comprehensive study significantly advances our understanding of drought regulatory atlas in plants and provides a robust foundation for future agricultural applications, particularly in the context of rapid environmental changes and the escalating challenges posed by global climate change.

## Material and Methods

### Drought Initiation and Nuclei Isolation

Approximately 2,000 seeds of *Arabidopsis thaliana* wild type accession Columbia (Col-0) were surface sterilized using a solution of 30% household bleach supplemented with 0.1% Triton X-100 (Thermo Scientific. Catalog number: A16046-AE) for 5 minutes. The seeds were then washed by 70% ethanol for 5 minutes and with sterile water for 5 times. The sterilized seeds were then diluted in 1 mL of 0.1% Agar solution. In a laminar flow hood, the seeds were pipetted onto square plates containing ½ solid Murashige and Skoog media containing 1% Sucrose and 0.8% Agar using 200ul pipette tips and evenly distributed alongside the horizontal grid. After plating, the plates were sealed with 3M™ Micropore™ Surgical Paper Tape (Fisher Scientific) and stored in the cold room for 48h. Subsequently, the plates were transferred to a growth chamber maintained at 23 °C with a 16h/8h light cycle and 50% humidity. Half of the plates were assigned to the drought group on day 6. Specifically, seedlings were transplanted onto PEG-infused ½ MS plates following the protocol by Paul E. Verslues for a 16-hour treatment. On day 7, seedlings from both the control and drought groups were collected for nuclei isolation. Freshly collected 7-day-old seedlings were subjected to nuclei extraction using the MinuteTM Single Nuclei Isolation Kit for Plant Tissues (Invent Biotechnology, Inc. PS-054) with necessary modifications to ensure optimal nuclei quality. Briefly, 400 mg of seedlings were immediately collected into a mortar containing 2 ml of buffer A. The seedlings were then ground using a pestle for approximately 2 min to facilitate the release of nuclei. The resulting slurry was filtered to a 50 mL tube using 40μm strainer (CORNING, catalog number: 07-201-430) and centrifuged at 2,000 g for 5 min. The pellet was dissolved in 500 μl of buffer A and then transferred to a 2 ml tube (CELLTREAT, catalog number: 229446) and centrifuged at 1,500 g for 5 min. The pellet was resuspended in 100 μl of buffer A plus 1 mL of buffer B by gently pipetting 20 times. The mixture was incubated on ice for 10 to remove most chloroplast. Then the mixture was centrifuged at 200 g for 2 min and the supernatant was transferred to a new 2 mL Tube (CELLTREAT, catalog number: 229446) and centrifuged at 1,000 g for 5 min. The pellet was dissolved in 500 μl of buffer C with 0.5% Triton X-100 and subjected to gentle shaking on an orbital shaker for 5 min. The suspension was then transferred to a filter cartridge provided by the kit in a 2 mL tube (CELLTREAT, catalog number: 229446), followed by centrifuging at 400 g for 2 min and 800 g for 4 min. The supernatant was completely removed, and the pellet was resuspended in 500 μl of 1X DPBS buffer (Gibco™. Catalog number: 14040141) with 1% BSA. The volume was then brought up to 2.5 mL into a 15 mL Falcon tube to ensure complete dilution of the nuclei. Prior to usage, all buffers and solutions mentioned above were freshly supplemented with 0.2% Protector RNase inhibitor (Millipore Sigma, catalog number: 3335402001).

### FACS and snMultiome library construction

After diluting the nuclei solution to 2.5 ml, 62.5 μL of BD 7-AAD Staining Solution (Fisher Scientific, catalog number: BDB559925) was added and mixed, followed by incubation on ice for approximately 10 minutes. The stained nuclei were then sorted using a BD FACSAria™ II flow cytometer (Flow Cytometry and Single Cell Core Facility, University of Alabama at Birmingham), with approximately 600,000 - 800,000 fluorescent events collected into a 1.5 ml microcentrifuge tube. The sorted nuclei were centrifuged at 800 g for 5 minutes, and the supernatant was carefully removed, leaving approximately 5 μL at the bottom. Next, 10 μL of 1X Diluted Nuclei Suspension Buffer (10X Genomics; PN-000207) supplemented with 0.2% Protector RNase Inhibitor and 1mM DTT was added to the nuclei solution, and gentle pipetting was performed 20 times to dissociate any nuclei clumps. Subsequently, 3 μL of the nuclei suspension was mixed with 9 μL of 0.4% Trypan Blue Solution (Thermo Fisher, catalog number: 15250061), and the nuclei count was determined under 20X brightfield microscope using a Chemglass Life Sciences Disposable Hemocytometer (Fisher Scientific, catalog number: 50-131-1352). The nuclei count was adjusted to 3000/μL using 1X Diluted Nuclei Suspension Buffer (10X Genomics; PN-000207). A total of 5 μL of the adjusted nuclei was used for Tn5 Tagmentation and subsequent library construction following the manufacturer’s instructions (10X Genomics, catalog number: CG000338). Subsequently, the constructed libraries, specifically designed for RNA sequencing and ATAC sequencing, were sequenced on the Illumina platform provided by the Genomics Core at the University of Alabama at Birmingham.

### Bioinformatic analysis

The sequenced reads were aligned to reference transcriptome and genome built by CellRanger-ARC. Obtained aligned reads for snRNA and snATAC were further input into Rstudio for filtering. snRNA and snATAC data were read into the Seurat object. Nuclei with RNA UMI more than 200 and less than 7000, gene counts more than 200, and mitochondrial reads less than 5% were filtered out. ATAC reads retained in filtered nuclei were further processed with peak calling using MACS2 with Arabidopsis-specific parameters. Post nuclei filtration, the RNA and ATAC reads in the cell matrix underwent normalization utilizing Seurat. The clustering of these datasets employed the FindClusters function in Seurat at a resolution of 0.5. Visualization was facilitated by the UMAP package. Integration of snRNA datasets from drought and water conditions utilized the Seurat V5 IntegrateData function (*56*). Cell identities were manually annotated based on a marker gene reference list compiled from various publications. Differential gene expression analysis between drought and control conditions was performed following the Seurat manual provided by the Satija lab (*56*). Pathway analysis for cluster marker genes was conducted using the gene set enrichment analysis (GSEA) package (*57*). Function analysis of differentially expressed genes in each cell type was undertaken using Metascape(*58*). Integration of RNA and ATAC data adhered to the methodologies outlined in published protocols (*32, 36*). Linkage analysis was done by employing the Singac function LinkPeaks and peaks specific to cell types are used for motif enrichment analysis. MINI-AC workflow from Manosalva et al. was applied for gene regulatory network inference (*39*). Pseudotime analysis of snRNA data was done with Monocle3 package (*59*), initiated from different cell types. Calculation of modules along the trajectory was performed within Monocle3, and genes enriched in each module were subject to functional analysis via Metascape.

## Author contributions

J.L. designed the experiments under the guidance of K.M.P.-M and M.S.M. A.M performed the majority of the bioinformatics analysis, while N. K. assisted in several bioinformatics analyses.

J.L. wrote the first draft and all authors edited the manuscript.

## Acknowledgments

This work was supported by NSF award IOS-2038872 to M.S.M. to K.M.P.-M. We thank the flow cytometry and the single-cell core facility teams at the University of Alabama at Birmingham for their assistance in library construction, and to the genomic core for their expertise in sequencing. Special thanks to Dr. Tatsuya Nobori for his technical guidance in single nuclei analysis and Dr. Wenbo Ma and Dr. Bozeng Tang for providing a cell marker reference list.

## Data and code availability

Upon request code will be provided.

## Ethics declarations

The authors declare no competing interests.

