## Supplemental Figures for "Single nuclei multiomics reveals the drought-driven gene regulatory atlas in Arabidopsis"

Fig. S1

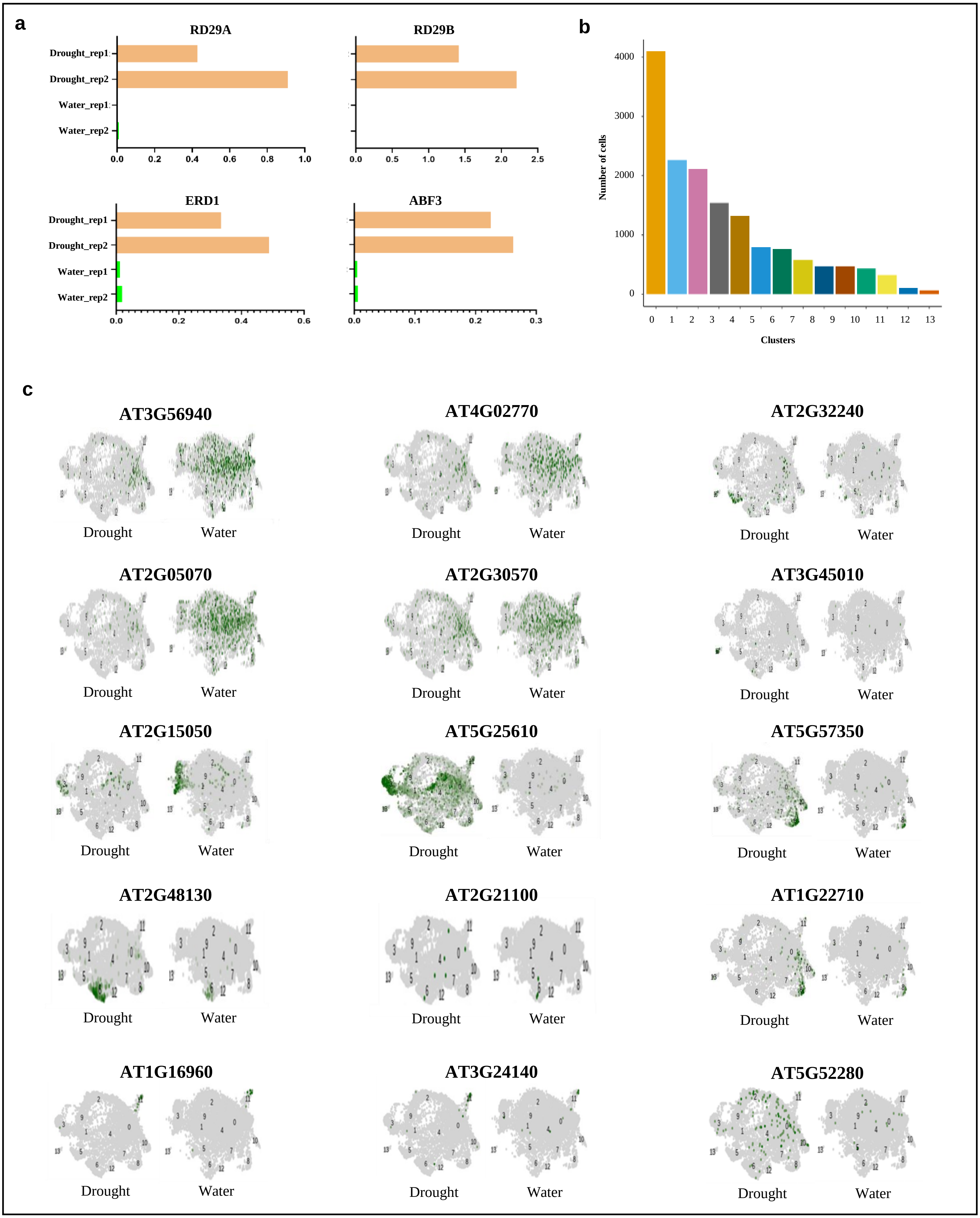
a
RD29A
RD29B
ERD1
ABF3
Drought_rep1
Drought_rep2
Water_rep1
Water_rep2
0.0 0.2 0.4 0.6 0.8 1.0
0.0 0.5 1.0 1.5 2.0 2.5
0.0 0.2 0.4 0.6
0.0 0.1 0.2 0.3
b
Number of cells
0 1000 2000 3000 4000
0 1 2 3 4 5 6 7 8 9 10 11 12 13
Clusters
c
AT3G56940
AT4G02770
AT2G32240
AT2G05070
AT2G30570
AT3G45010
AT2G15050
AT5G25610
AT5G57350
AT2G48130
AT2G21100
AT1G22710
AT1G16960
AT3G24140
AT5G52280
Drought
Water

**Fig. S1: Construction of drought-simulated single nuclei transcriptomic atlas**

(A) Transcript quantification of classic drought responsive genes using Real-Time Quantitative Reverse Transcription PCR (qRT-PCR) in drought simulated and control groups.
(B) Nuclei counting across snRNA clusters.
(C) Uniform Manifold Approximation and Projection (UMAP) inlet of collected cell type markers.

Fig. S2

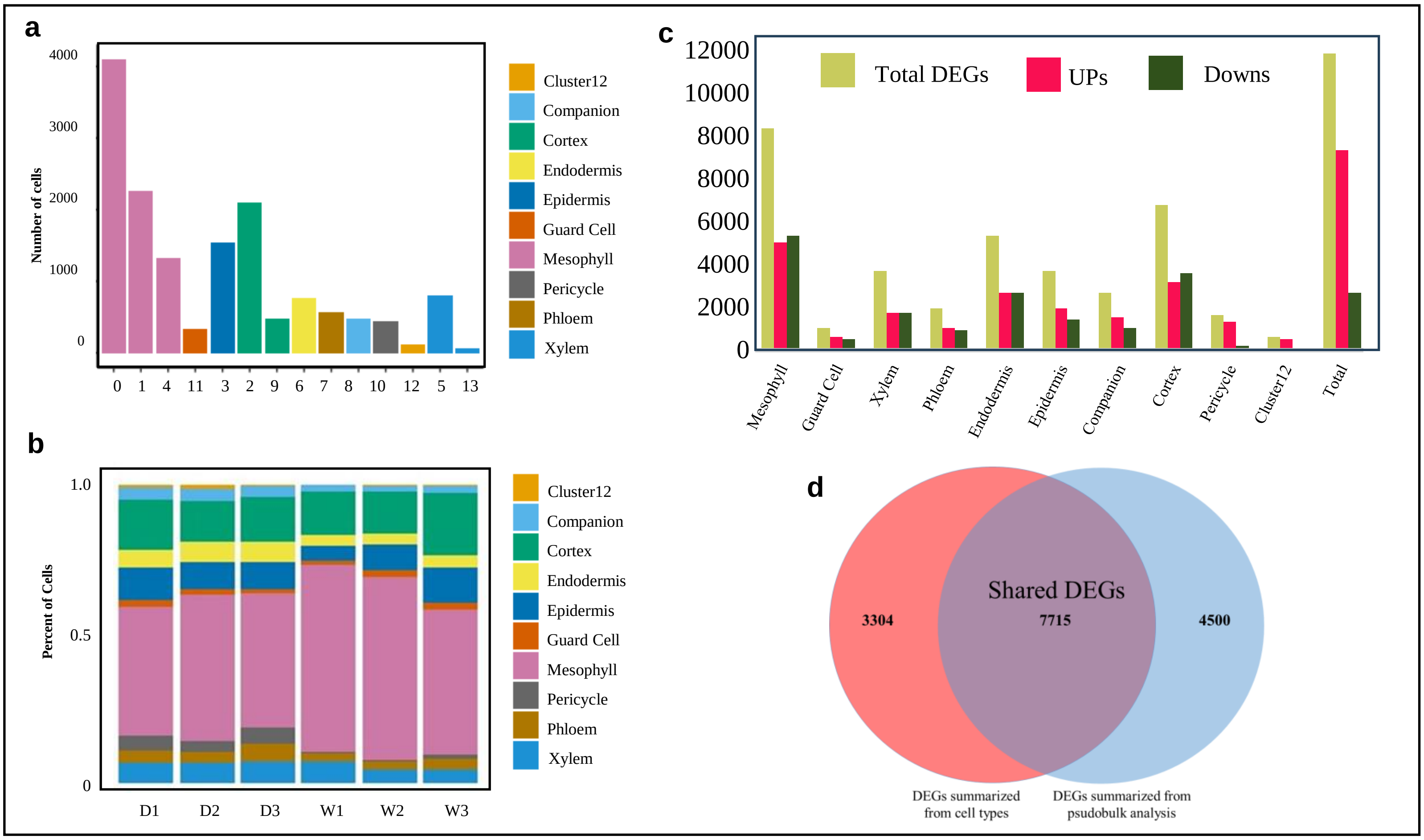

a
4000
3000
2000
1000
0
Number of cells
0 1 4 11 3 2 9 6 7 8 10 12 5 13
Cluster12
Companion
Cortex
Endodermis
Epidermis
Guard Cell
Mesophyll
Pericycle
Phloem
Xylem
b
1.0
0.5
0
Percent of Cells
D1 D2 D3 W1 W2 W3
Cluster12
Companion
Cortex
Endodermis
Epidermis
Guard Cell
Mesophyll
Pericycle
Phloem
Xylem
c
12000
10000
8000
8000
6000
4000
2000
0
Total DEGs
UPs
Downs
Mesophyll
Guard Cell
Xylem
Phloem
Endodermis
Epidermis
Companion
Cortex
Pericycle
Cluster12
Total
d
Shared DEGs
3304
7715
4500
DEGs summarized
from cell types
DEGs summarized from
psudobulk analysis

**Fig. S2: Single nuclei transcriptomic analysis assign major cell types of roots and shoots.**

(A) Number of nuclei across cell types and clusters.
(B) Percent of cell types exhibited by each replicate of drought and water samples.
(C) Number of Differentially expressed genes (DEGs) in each cell types and whole tissue.
(D) Venn diagram showing shared and unique DEGs to single cell and pseudobulk analysis.

**Fig. S3.**

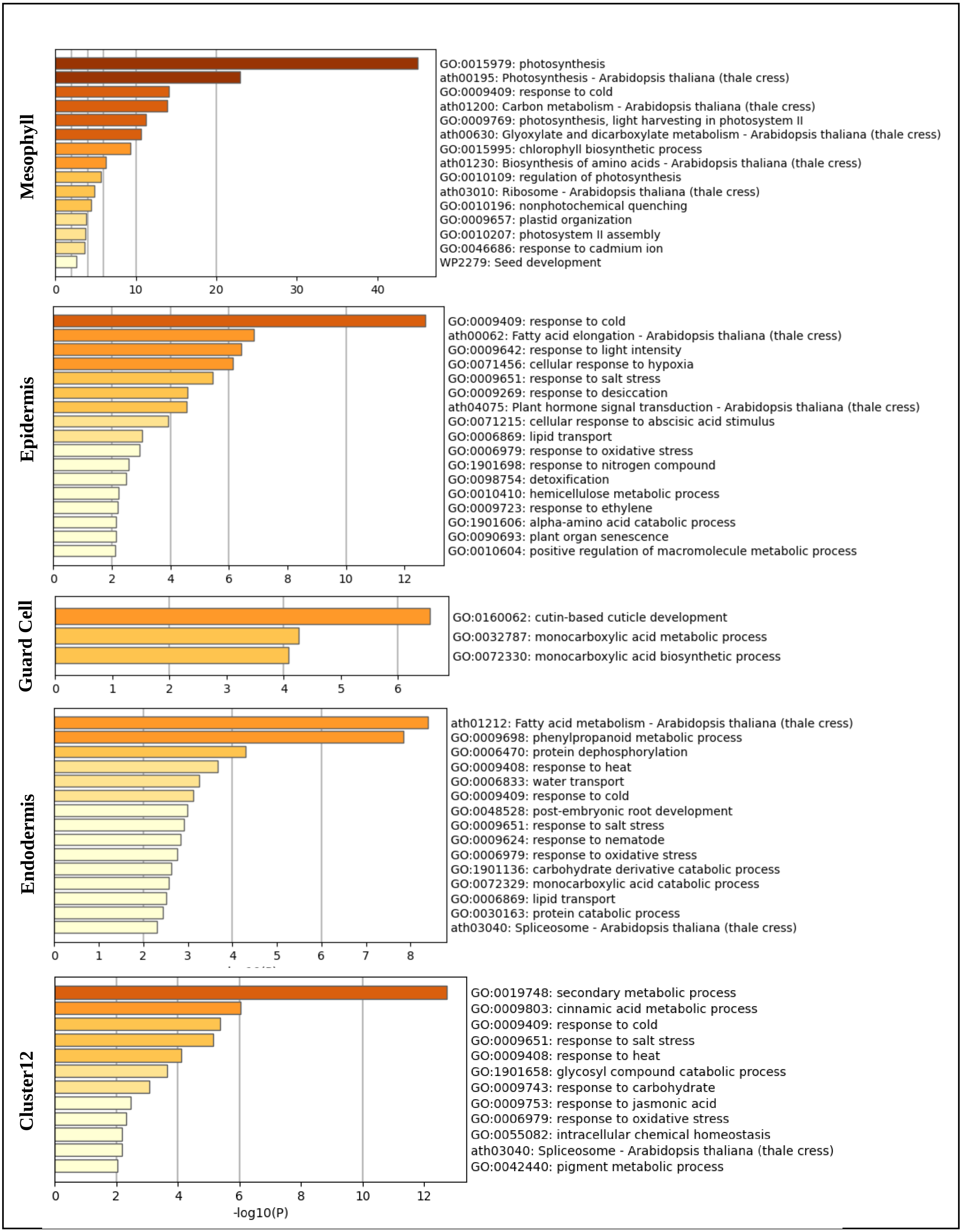
Mesophyll
GO:0015979: photosynthesis
ath00195: Photosynthesis - Arabidopsis thaliana (thale cress)
GO:0009409: response to cold
ath01200: Carbon metabolism - Arabidopsis thaliana (thale cress)
GO:0009769: photosynthesis, light harvesting in photosystem II
ath00630: Glyoxylate and dicarboxylate metabolism - Arabidopsis thaliana (thale cress)
GO:0015995: chlorophyll biosynthetic process
ath01230: Biosynthesis of amino acids - Arabidopsis thaliana (thale cress)
GO:0010109: regulation of photosynthesis
ath03010: Ribosome - Arabidopsis thaliana (thale cress)
GO:0010196: nonphotochemical quenching
GO:0009657: plastid organization
GO:0010207: photosystem II assembly
GO:0046686: response to cadmium ion
WP2279: Seed development
0 10 20 30 40
Epidermis
GO:0009409: response to cold
ath00062: Fatty acid elongation - Arabidopsis thaliana (thale cress)
GO:0009642: response to light intensity
GO:0071456: cellular response to hypoxia
GO:0009651: response to salt stress
GO:0009269: response to desiccation
ath04075: Plant hormone signal transduction - Arabidopsis thaliana (thale cress)
GO:0071215: cellular response to abscisic acid stimulus
GO:0006869: lipid transport
GO:0006979: response to oxidative stress
GO:1901698: response to nitrogen compound
GO:0098754: detoxification
GO:0010410: hemicellulose metabolic process
GO:0009723: response to ethylene
GO:1901606: alpha-amino acid catabolic process
GO:0090693: plant organ senescence
GO:0010604: positive regulation of macromolecule metabolic process
0 2 4 6 8 10 12
Guard Cell
GO:0160062: cutin-based cuticle development
GO:0032787: monocarboxylic acid metabolic process
GO:0072330: monocarboxylic acid biosynthetic process
0 1 2 3 4 5 6
Endodermis
ath01212: Fatty acid metabolism - Arabidopsis thaliana (thale cress)
GO:0009698: phenylpropanoid metabolic process
GO:0006470: protein dephosphorylation
GO:0009408: response to heat
GO:0006833: water transport
GO:0009409: response to cold
GO:0048528: post-embryonic root development
GO:0009651: response to salt stress
GO:0009624: response to nematode
GO:0006979: response to oxidative stress
GO:1901136: carbohydrate derivative catabolic process
GO:0072329: monocarboxylic acid catabolic process
GO:0006869: lipid transport
GO:0030163: protein catabolic process
ath03040: Spliceosome - Arabidopsis thaliana (thale cress)
0 1 2 3 4 5 6 7 8
Cluster12
GO:0019748: secondary metabolic process
GO:0009803: cinnamic acid metabolic process
GO:0009409: response to cold
GO:0009651: response to salt stress
GO:0009408: response to heat
GO:1901658: glycosyl compound catabolic process
GO:0009743: response to carbohydrate
GO:0009753: response to jasmonic acid
GO:0006979: response to oxidative stress
GO:0055082: intracellular chemical homeostasis
ath03040: Spliceosome - Arabidopsis thaliana (thale cress)
GO:0042440: pigment metabolic process
0 2 4 6 8 10 12
-log10(P)

**Fig. S3 (Continued)**

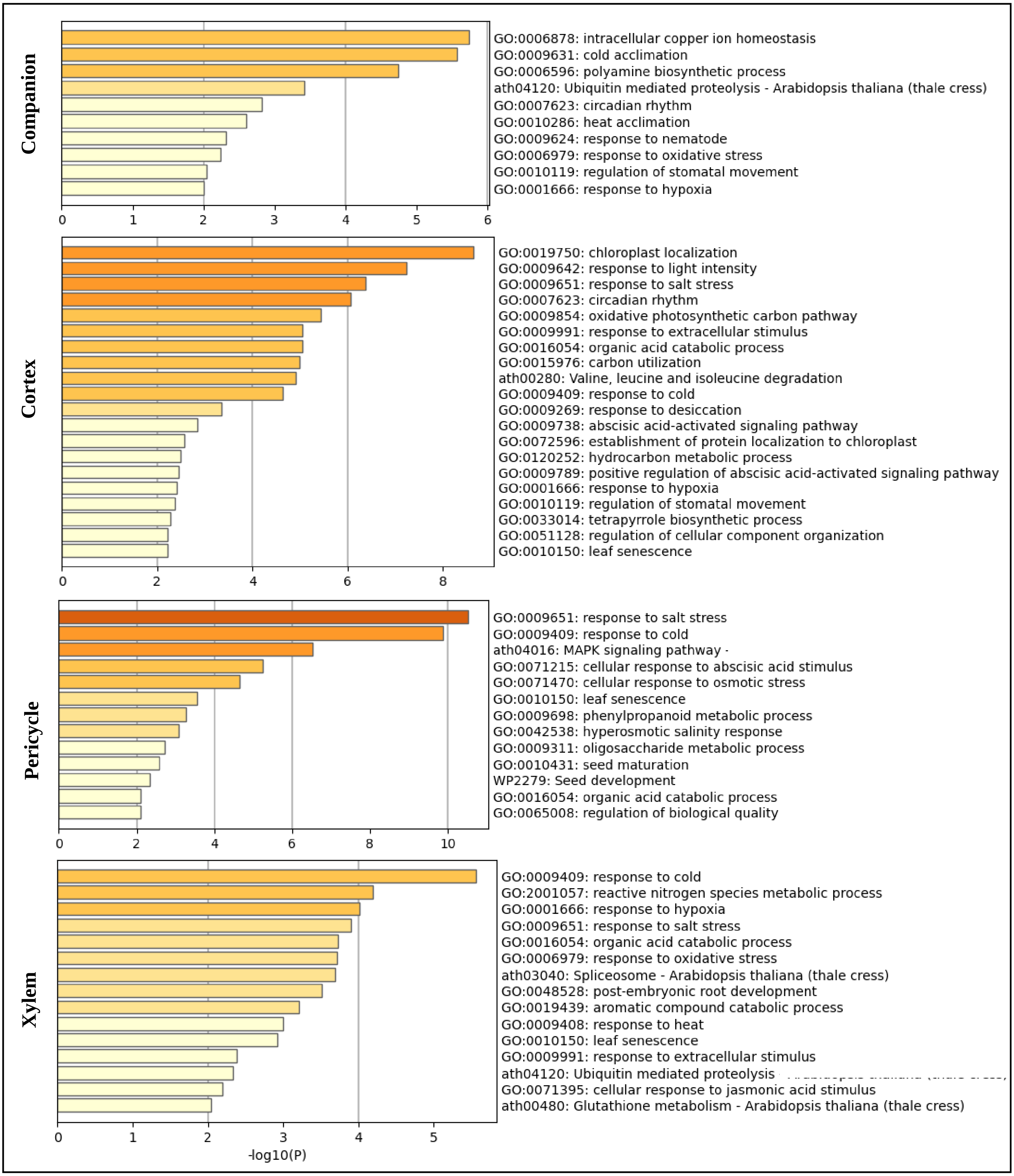

**Fig. S3: Pathway analysis of differential expressed genes present in each cell type.**

**Fig. S4**

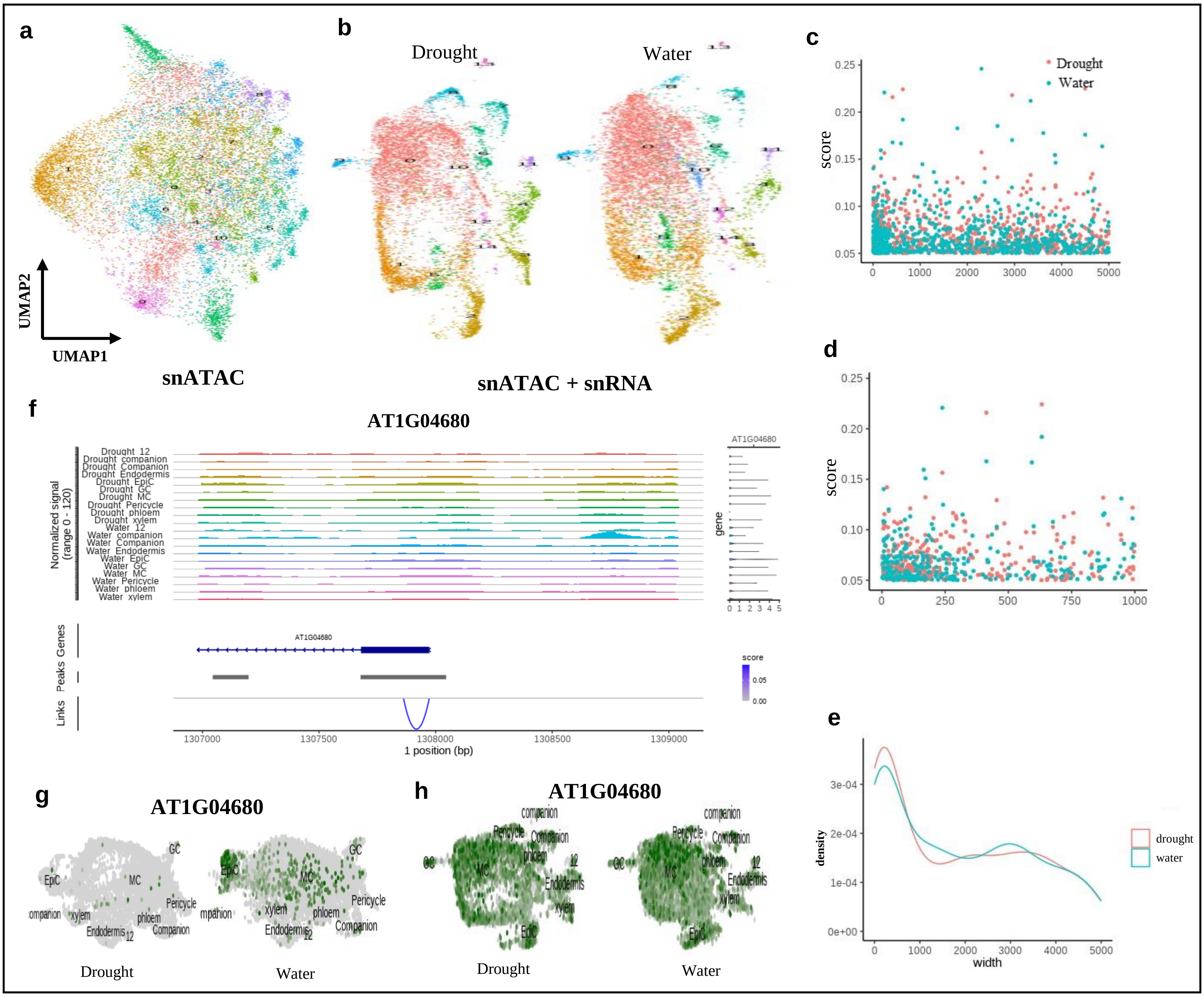

a
b
c
Drought
Water
UMAP2
UMAP1
snATAC
snATAC + snRNA
score
d
f
AT1G04680
Normalized signal (range 0 - 120)
Drought_12
Drought_companion
Drought_Companion
Drought_Endodermis
Drought_EpiC
Drought_GC
Drought_MC
Drought_Pericycle
Drought_phloem
Drought_xylem
Water_12
Water_companion
Water_Companion
Water_Endodermis
Water_EpiC
Water_GC
Water_MC
Water_Pericycle
Water_phloem
Water_xylem
gene
Genes
Peaks
Links
score
1 position (bp)
e
density
width
drought
water
g
h
EpiC
MC
GC
Pericycle
phloem
xylem
Endodermis
Companion
companion
12
Drought
Water

**Fig. S4: Linkage between single nuclei RNA (snRNA) and ATAC (snATAC) data allows to identify cell type specific regulators.**

(A) Uniform Manifold Approximation and Projection (UMAP) plots of single nuclei ATAC data. Nuclei are colored by Leiden clusters.
(B) UMAP plots of embedding snRNA and snATAC data based on treatment. Nuclei are colored by Leiden clusters.
(C) Scatter plot indicating the linkage score distribution along 5000bp upstream and downstream of transcriptional start site (TSS)
(D) Scatter plot indicating the linkage score distribution along 1000bp upstream and downstream of transcriptional start site (TSS)
(E) Scatter plot indicating the linkage score distribution along 600bp upstream and downstream of transcriptional start site (TSS)
(F) Chromatin coverage plots showing cluster-specific linkage, exemplified by epidermis marker gene AT1G04680.
(G) Uniform Manifold Approximation and Projection (UMAP) inlet of AT1G04680 in snRNA data of drought and water groups.
(H) Uniform Manifold Approximation and Projection (UMAP) inlet of AT1G04680 in integrated snRNA and snATAC data of drought and water samples.

Fig. S5

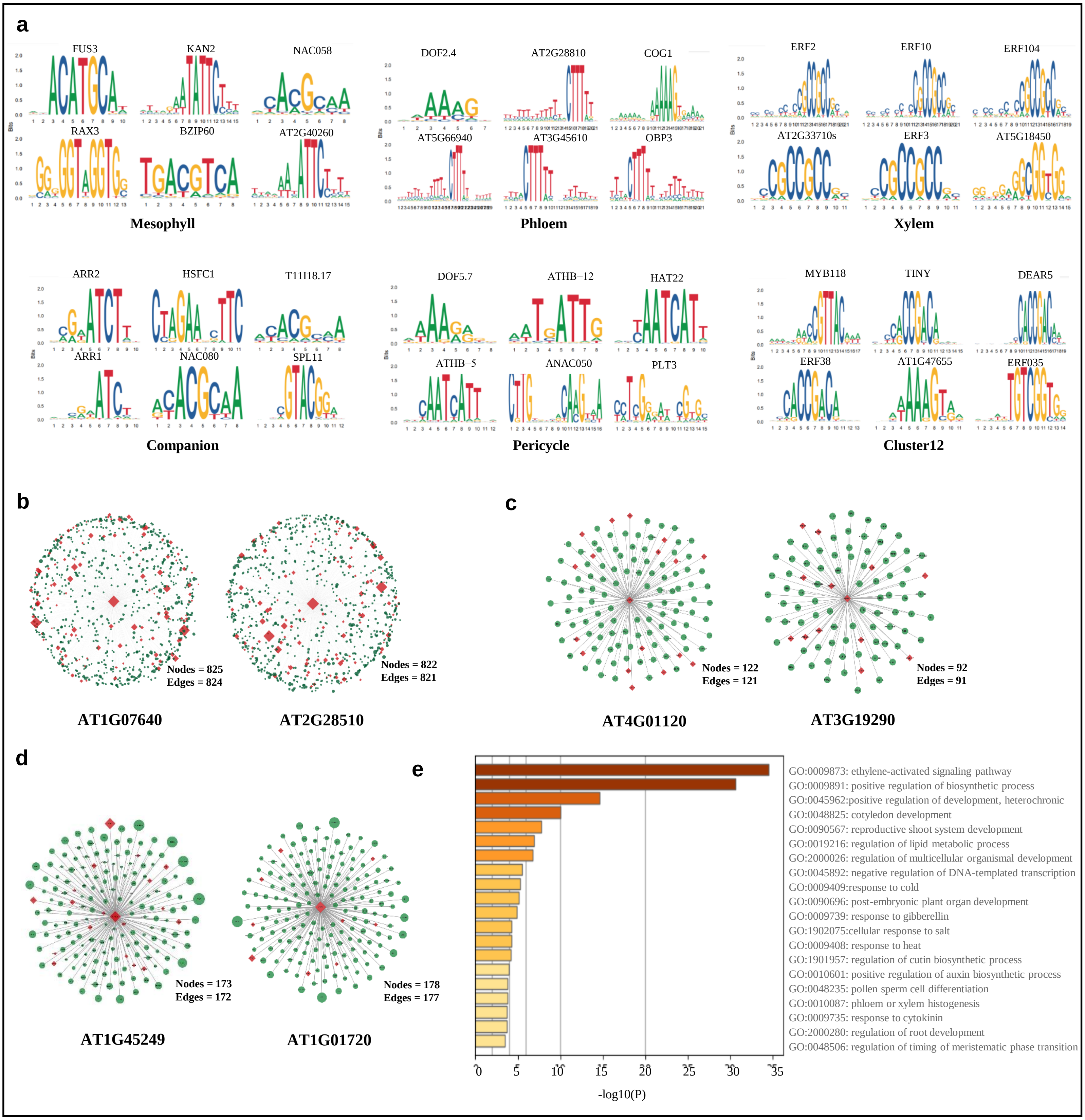
a
FUS3
KAN2
NAC058
RAX3
BZIP60
AT2G40260
Mesophyll
DOF2.4
AT2G28810
COG1
AT5G66940
AT3G45610
OBP3
Phloem
ERF2
ERF10
ERF104
AT2G33710s
ERF3
AT5G18450
Xylem
ARR2
HSFC1
T11I18.17
ARR1
NAC080
SPL11
Companion
DOF5.7
ATHB−12
HAT22
ATHB−5
ANAC050
PLT3
Pericycle
MYB118
TINY
DEAR5
ERF38
AT1G47655
ERF035
Cluster12
b
Nodes = 825
Edges = 824
AT1G07640
Nodes = 822
Edges = 821
AT2G28510
c
Nodes = 122
Edges = 121
AT4G01120
Nodes = 92
Edges = 91
AT3G19290
d
Nodes = 173
Edges = 172
AT1G45249
Nodes = 178
Edges = 177
AT1G01720
e
GO:0009873: ethylene-activated signaling pathway
GO:0009891: positive regulation of biosynthetic process
GO:0045962:positive regulation of development, heterochronic
GO:0048825: cotyledon development
GO:0090567: reproductive shoot system development
GO:0019216: regulation of lipid metabolic process
GO:2000026: regulation of multicellular organismal development
GO:0045892: negative regulation of DNA-templated transcription
GO:0009409:response to cold
GO:0090696: post-embryonic plant organ development
GO:0009739: response to gibberellin
GO:1902075:cellular response to salt
GO:0009408: response to heat
GO:1901957: regulation of cutin biosynthetic process
GO:0010601: positive regulation of auxin biosynthetic process
GO:0048235: pollen sperm cell differentiation
GO:0010087: phloem or xylem histogenesis
GO:0009735: response to cytokinin
GO:2000280: regulation of root development
GO:0048506: regulation of timing of meristematic phase transition
0
5
10
15
20
25
30
35
-log10(P)

**Fig. S5: Linking transcriptomic profiles and chromatin accessibility identified cell type-specific and condition-specific regulatory events.**

(A) Top motif enriched in mesophyll, phloem, xylem, companion, pericycle and cluster12. Motif is inferred by motif enrichment analysis.

(B) Gene regulatory network (GRN) inference of top transcriptional factors marked by MINI-AC analysis. Exemplified by epidermis marker, AT1G07640 and AT2G28510.

(C) GRN inference of top transcriptional factors marked by MINI-AC analysis. Exemplified by endodermis marker, AT4G01120 and AT3G19290.

(D) GRN inference of top transcriptional factors marked by MINI-AC analysis. Exemplified by epidermis marker, AT1G45249 and AT1G01720.

(E) Pathway analysis of GRN network inferred by MINI-AC in mesophyll.

Fig. S6

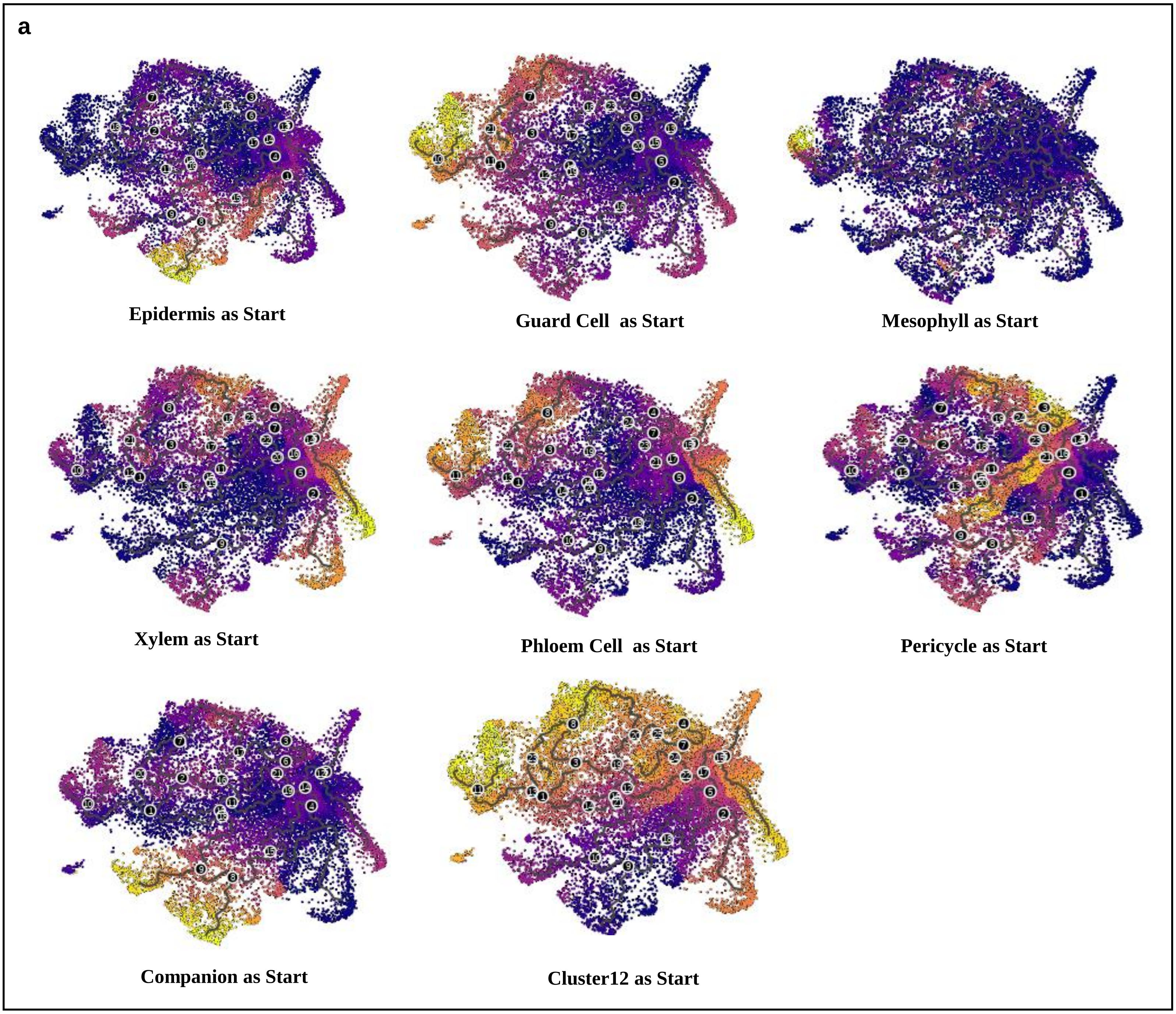
a
Epidermis as Start
Guard Cell as Start
Mesophyll as Start
Xylem as Start
Phloem Cell as Start
Pericycle as Start
Companion as Start
Cluster12 as Start

**Fig. S6: Pseduotime analysis of all cell types indicates different transcriptional progress patterns using distinct starting point.**

Fig. S7

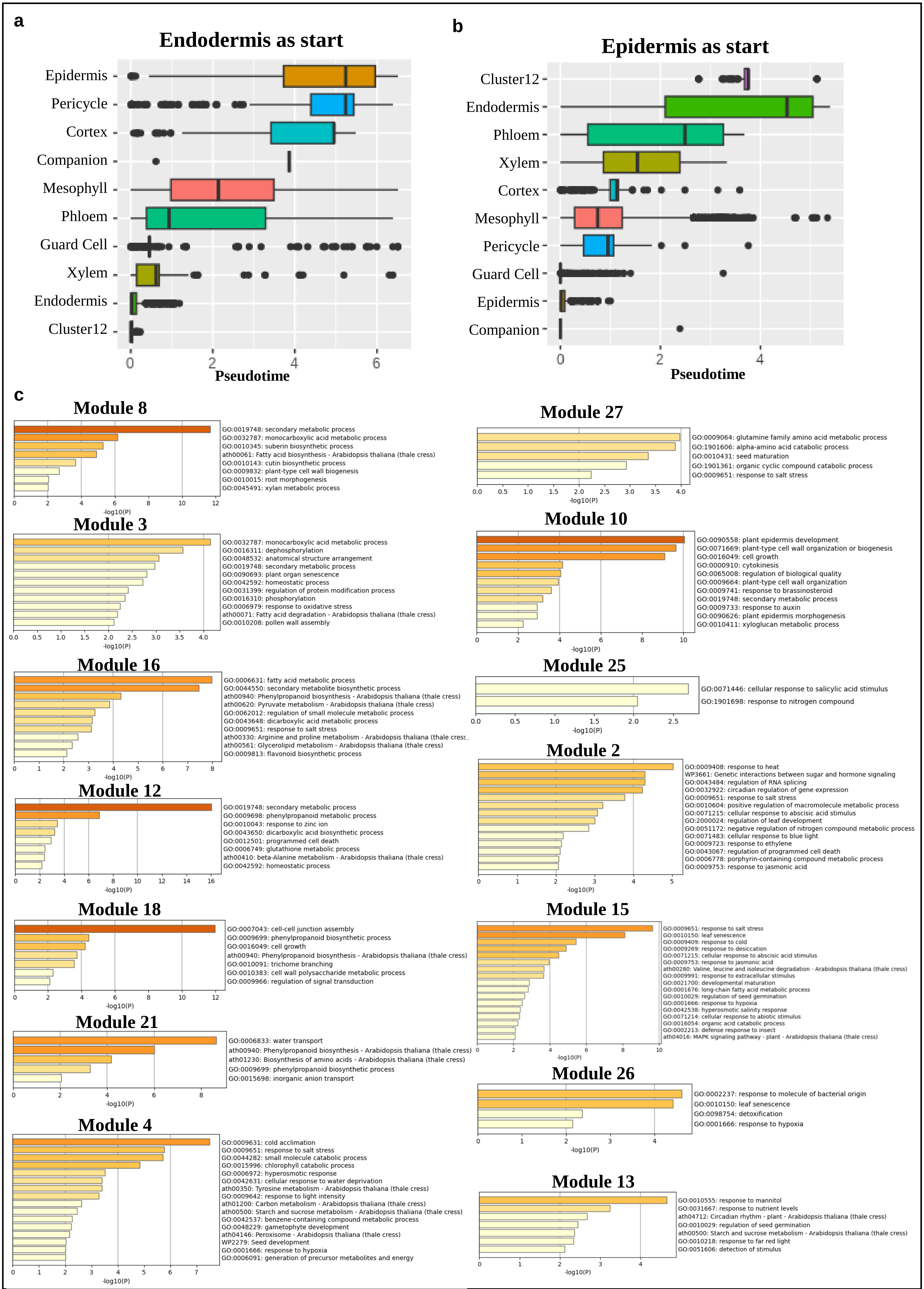
a
Endodermis as start
Epidermis
Pericycle
Cortex
Companion
Mesophyll
Phloem
Guard Cell
Xylem
Endodermis
Cluster12
Pseudotime
b
Epidermis as start
Cluster12
Endodermis
Phloem
Xylem
Cortex
Mesophyll
Pericycle
Guard Cell
Epidermis
Companion
Pseudotime
c
Module 8
GO:0019748: secondary metabolic process
GO:0032787: monocarboxylic acid metabolic process
GO:0010345: suberin biosynthetic process
ath00061: Fatty acid biosynthesis - Arabidopsis thaliana (thale cress)
GO:0010143: cutin biosynthetic process
GO:0009832: plant-type cell wall biogenesis
GO:0010015: root morphogenesis
GO:0045491: xylan metabolic process
-log10(P)
Module 3
GO:0032787: monocarboxylic acid metabolic process
GO:0016311: dephosphorylation
GO:0048532: anatomical structure arrangement
GO:0019748: secondary metabolic process
GO:0090693: plant organ senescence
GO:0042592: homeostatic process
GO:0031399: regulation of protein modification process
GO:0016310: phosphorylation
GO:0006979: response to oxidative stress
ath00071: Fatty acid degradation - Arabidopsis thaliana (thale cress)
GO:0010208: pollen wall assembly
-log10(P)
Module 16
GO:0006631: fatty acid metabolic process
GO:0044550: secondary metabolite biosynthetic process
ath00940: Phenylpropanoid biosynthesis - Arabidopsis thaliana (thale cress)
ath00620: Pyruvate metabolism - Arabidopsis thaliana (thale cress)
GO:0062012: regulation of small molecule metabolic process
GO:0043648: dicarboxylic acid metabolic process
GO:0009651: response to salt stress
ath00330: Arginine and proline metabolism - Arabidopsis thaliana (thale cress
ath00561: Glycerolipid metabolism - Arabidopsis thaliana (thale cress)
GO:0009813: flavonoid biosynthetic process
-log10(P)
Module 12
GO:0019748: secondary metabolic process
GO:0009698: phenylpropanoid metabolic process
GO:0010043: response to zinc ion
GO:0043650: dicarboxylic acid biosynthetic process
GO:0012501: programmed cell death
GO:0006749: glutathione metabolic process
ath00410: beta-Alanine metabolism - Arabidopsis thaliana (thale cress)
GO:0042592: homeostatic process
-log10(P)
Module 18
GO:0007043: cell-cell junction assembly
GO:0009699: phenylpropanoid biosynthetic process
GO:0016049: cell growth
ath00940: Phenylpropanoid biosynthesis - Arabidopsis thaliana (thale cress)
GO:0010091: trichome branching
GO:0010383: cell wall polysaccharide metabolic process
GO:0009966: regulation of signal transduction
-log10(P)
Module 21
GO:0006833: water transport
ath00940: Phenylpropanoid biosynthesis - Arabidopsis thaliana (thale cress)
ath01230: Biosynthesis of amino acids - Arabidopsis thaliana (thale cress)
GO:0009699: phenylpropanoid biosynthetic process
GO:0015698: inorganic anion transport
-log10(P)
Module 4
GO:0009631: cold acclimation
GO:0009651: response to salt stress
GO:0044282: small molecule catabolic process
GO:0015996: chlorophyll catabolic process
GO:0006972: hyperosmotic response
GO:0042631: cellular response to water deprivation
ath00350: Tyrosine metabolism - Arabidopsis thaliana (thale cress)
GO:0009642: response to light intensity
ath01200: Carbon metabolism - Arabidopsis thaliana (thale cress)
ath00500: Starch and sucrose metabolism - Arabidopsis thaliana (thale cress)
GO:0042537: benzene-containing compound metabolic process
GO:0048229: gametophyte development
ath04146: Peroxisome - Arabidopsis thaliana (thale cress)
WP2279: Seed development
GO:0001666: response to hypoxia
GO:0006091: generation of precursor metabolites and energy
-log10(P)
Module 27
GO:0009064: glutamine family amino acid metabolic process
GO:1901606: alpha-amino acid catabolic process
GO:0010431: seed maturation
GO:1901361: organic cyclic compound catabolic process
GO:0009651: response to salt stress
-log10(P)
Module 10
GO:0090558: plant epidermis development
GO:0071669: plant-type cell wall organization or biogenesis
GO:0016049: cell growth
GO:0000910: cytokinesis
GO:0065008: regulation of biological quality
GO:0009664: plant-type cell wall organization
GO:0009741: response to brassinosteroid
GO:0019748: secondary metabolic process
GO:0009733: response to auxin
GO:0090626: plant epidermis morphogenesis
GO:0010411: xyloglucan metabolic process
-log10(P)
Module 25
GO:0071446: cellular response to salicylic acid stimulus
GO:1901698: response to nitrogen compound
-log10(P)
Module 2
GO:0009408: response to heat
WP3661: Genetic interactions between sugar and hormone signaling
GO:0043484: regulation of RNA splicing
GO:0032922: circadian regulation of gene expression
GO:0009651: response to salt stress
GO:0010604: positive regulation of macromolecule metabolic process
GO:0071215: cellular response to abscisic acid stimulus
GO:2000024: regulation of leaf development
GO:0051172: negative regulation of nitrogen compound metabolic process
GO:0071483: cellular response to blue light
GO:0009723: response to ethylene
GO:0043067: regulation of programmed cell death
GO:0006778: porphyrin-containing compound metabolic process
GO:0009753: response to jasmonic acid
-log10(P)
Module 15
GO:0009651: response to salt stress
GO:0010150: leaf senescence
GO:0009409: response to cold
GO:0009269: response to desiccation
GO:0071215: cellular response to abscisic acid stimulus
GO:0009753: response to jasmonic acid
ath00280: Valine, leucine and isoleucine degradation - Arabidopsis thaliana (thale cress)
GO:0009991: response to extracellular stimulus
GO:0021700: developmental maturation
GO:0001676: long-chain fatty acid metabolic process
GO:0010029: regulation of seed germination
GO:0001666: response to hypoxia
GO:0042538: hyperosmotic salinity response
GO:0071214: cellular response to abiotic stimulus
GO:0016054: organic acid catabolic process
GO:0002213: defense response to insect
ath04016: MAPK signaling pathway - plant - Arabidopsis thaliana (thale cress)
-log10(P)
Module 26
GO:0002237: response to molecule of bacterial origin
GO:0010150: leaf senescence
GO:0098754: detoxification
GO:0001666: response to hypoxia
-log10(P)
Module 13
GO:0010555: response to mannitol
GO:0031667: response to nutrient levels
ath04712: Circadian rhythm - plant - Arabidopsis thaliana (thale cress)
GO:0010029: regulation of seed germination
ath00500: Starch and sucrose metabolism - Arabidopsis thaliana (thale cress)
GO:0010218: response to far red light
GO:0051606: detection of stimulus
-log10(P)

**Fig. S7**

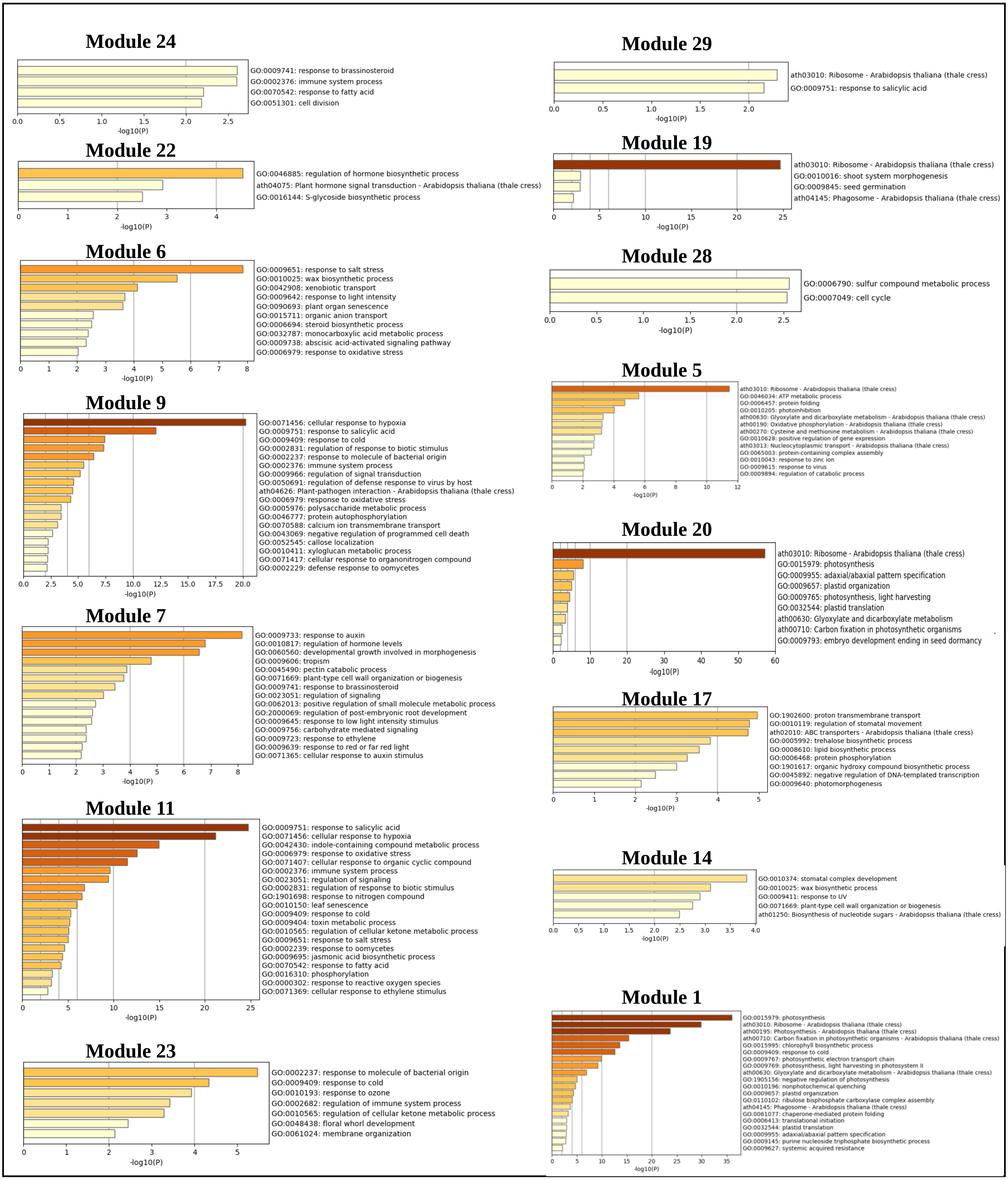
Module 24
GO:0009741: response to brassinosteroid
GO:0002376: immune system process
GO:0070542: response to fatty acid
GO:0051301: cell division
0.0 0.5 1.0 1.5 2.0 2.5
-log10(P)
Module 22
GO:0046885: regulation of hormone biosynthetic process
ath04075: Plant hormone signal transduction - Arabidopsis thaliana (thale cress)
GO:0016144: S-glycoside biosynthetic process
0 1 2 3 4
-log10(P)
Module 6
GO:0009651: response to salt stress
GO:0010025: wax biosynthetic process
GO:0042908: xenobiotic transport
GO:0009642: response to light intensity
GO:0090693: plant organ senescence
GO:0015711: organic anion transport
GO:0006694: steroid biosynthetic process
GO:0032787: monocarboxylic acid metabolic process
GO:0009738: abscisic acid-activated signaling pathway
GO:0006979: response to oxidative stress
0 1 2 3 4 5 6 7 8
-log10(P)
Module 9
GO:0071456: cellular response to hypoxia
GO:0009751: response to salicylic acid
GO:0009409: response to cold
GO:0002831: regulation of response to biotic stimulus
GO:0002237: response to molecule of bacterial origin
GO:0002376: immune system process
GO:0009966: regulation of signal transduction
GO:0050691: regulation of defense response to virus by host
ath04626: Plant-pathogen interaction - Arabidopsis thaliana (thale cress)
GO:0006979: response to oxidative stress
GO:0005976: polysaccharide metabolic process
GO:0046777: protein autophosphorylation
GO:0070588: calcium ion transmembrane transport
GO:0043069: negative regulation of programmed cell death
GO:0052545: callose localization
GO:0010411: xyloglucan metabolic process
GO:0071417: cellular response to organonitrogen compound
GO:0002229: defense response to oomycetes
0.0 2.5 5.0 7.5 10.0 12.5 15.0 17.5 20.0
-log10(P)
Module 7
GO:0009733: response to auxin
GO:0010817: regulation of hormone levels
GO:0060560: developmental growth involved in morphogenesis
GO:0009606: tropism
GO:0045490: pectin catabolic process
GO:0071669: plant-type cell wall organization or biogenesis
GO:0009741: response to brassinosteroid
GO:0023051: regulation of signaling
GO:0062013: positive regulation of small molecule metabolic process
GO:2000069: regulation of post-embryonic root development
GO:0009645: response to low light intensity stimulus
GO:0009756: carbohydrate mediated signaling
GO:0009723: response to ethylene
GO:0009639: response to red or far red light
GO:0071365: cellular response to auxin stimulus
0 1 2 3 4 5 6 7 8
-log10(P)
Module 11
GO:0009751: response to salicylic acid
GO:0071456: cellular response to hypoxia
GO:0042430: indole-containing compound metabolic process
GO:0006979: response to oxidative stress
GO:0071407: cellular response to organic cyclic compound
GO:0002376: immune system process
GO:0023051: regulation of signaling
GO:0002831: regulation of response to biotic stimulus
GO:1901698: response to nitrogen compound
GO:0010150: leaf senescence
GO:0009409: response to cold
GO:0009404: toxin metabolic process
GO:0010565: regulation of cellular ketone metabolic process
GO:0009651: response to salt stress
GO:0002239: response to oomycetes
GO:0009695: jasmonic acid biosynthetic process
GO:0070542: response to fatty acid
GO:0016310: phosphorylation
GO:0000302: response to reactive oxygen species
GO:0071369: cellular response to ethylene stimulus
0 5 10 15 20 25
-log10(P)
Module 23
GO:0002237: response to molecule of bacterial origin
GO:0009409: response to cold
GO:0010193: response to ozone
GO:0002682: regulation of immune system process
GO:0010565: regulation of cellular ketone metabolic process
GO:0048438: floral whorl development
GO:0061024: membrane organization
0 1 2 3 4 5
-log10(P)
Module 29
ath03010: Ribosome - Arabidopsis thaliana (thale cress)
GO:0009751: response to salicylic acid
0.0 0.5 1.0 1.5 2.0
-log10(P)
Module 19
ath03010: Ribosome - Arabidopsis thaliana (thale cress)
GO:0010016: shoot system morphogenesis
GO:0009845: seed germination
ath04145: Phagosome - Arabidopsis thaliana (thale cress)
0 5 10 15 20 25
-log10(P)
Module 28
GO:0006790: sulfur compound metabolic process
GO:0007049: cell cycle
0.0 0.5 1.0 1.5 2.0 2.5
-log10(P)
Module 5
ath03010: Ribosome - Arabidopsis thaliana (thale cress)
GO:0046034: ATP metabolic process
GO:0006457: protein folding
GO:0010205: photoinhibition
ath00630: Glyoxylate and dicarboxylate metabolism - Arabidopsis thaliana (thale cress)
ath00190: Oxidative phosphorylation - Arabidopsis thaliana (thale cress)
ath00270: Cysteine and methionine metabolism - Arabidopsis thaliana (thale cress)
GO:0010628: positive regulation of gene expression
ath03013: Nucleocytoplasmic transport - Arabidopsis thaliana (thale cress)
GO:0065003: protein-containing complex assembly
GO:0010043: response to zinc ion
GO:0009615: response to virus
GO:0009894: regulation of catabolic process
0 2 4 6 8 10 12
-log10(P)
Module 20
ath03010: Ribosome - Arabidopsis thaliana (thale cress)
GO:0015979: photosynthesis
GO:0009955: adaxial/abaxial pattern specification
GO:0009657: plastid organization
GO:0009765: photosynthesis, light harvesting
GO:0032544: plastid translation
ath00630: Glyoxylate and dicarboxylate metabolism
ath00710: Carbon fixation in photosynthetic organisms
GO:0009793: embryo development ending in seed dormancy
0 10 20 30 40 50 60
-log10(P)
Module 17
GO:1902600: proton transmembrane transport
GO:0010119: regulation of stomatal movement
ath02010: ABC transporters - Arabidopsis thaliana (thale cress)
GO:0005992: trehalose biosynthetic process
GO:0008610: lipid biosynthetic process
GO:0006468: protein phosphorylation
GO:1901617: organic hydroxy compound biosynthetic process
GO:0045892: negative regulation of DNA-templated transcription
GO:0009640: photomorphogenesis
0 1 2 3 4 5
-log10(P)
Module 14
GO:0010374: stomatal complex development
GO:0010025: wax biosynthetic process
GO:0009411: response to UV
GO:0071669: plant-type cell wall organization or biogenesis
ath01250: Biosynthesis of nucleotide sugars - Arabidopsis thaliana (thale cress)
0.0 0.5 1.0 1.5 2.0 2.5 3.0 3.5 4.0
-log10(P)
Module 1
GO:0015979: photosynthesis
ath03010: Ribosome - Arabidopsis thaliana (thale cress)
ath00195: Photosynthesis - Arabidopsis thaliana (thale cress)
ath00710: Carbon fixation in photosynthetic organisms - Arabidopsis thaliana (thale cress)
GO:0015995: chlorophyll biosynthetic process
GO:0009409: response to cold
GO:0009767: photosynthetic electron transport chain
GO:0009769: photosynthesis, light harvesting in photosystem II
ath00630: Glyoxylate and dicarboxylate metabolism - Arabidopsis thaliana (thale cress)
GO:1905156: negative regulation of photosynthesis
GO:0010196: nonphotochemical quenching
GO:0009657: plastid organization
GO:0110102: ribulose bisphosphate carboxylase complex assembly
ath04145: Phagosome - Arabidopsis thaliana (thale cress)
GO:0061077: chaperone-mediated protein folding
GO:0006413: translational initiation
GO:0032544: plastid translation
GO:0009955: adaxial/abaxial pattern specification
GO:0009145: purine nucleoside triphosphate biosynthetic process
GO:0009627: systemic acquired resistance
0 5 10 15 20 25 30 35
-log10(P)

**Fig. S7: Pathway analysis of monocle3-inferred modules reveals the sequential transcriptional events in endodermis, epidermis and guard cell.**

(A) Boxplot showing the transcriptional dynamics across cell types, with endodermis as start point.

(B) Boxplot showing the transcriptional dynamics across cell types, with epidermis as start point.

(C) Pathway analysis of distinct modules exhibited by endodermis, epidermis and guard cell.
